# Stabilizing brain-computer interfaces through alignment of latent dynamics

**DOI:** 10.1101/2022.04.06.487388

**Authors:** Brianna M. Karpowicz, Yahia H. Ali, Lahiru N. Wimalasena, Andrew R. Sedler, Mohammad Reza Keshtkaran, Kevin Bodkin, Xuan Ma, Lee E. Miller, Chethan Pandarinath

**Affiliations:** Wallace H. Coulter Department of Biomedical Engineering, Emory University and Georgia Institute of Technology, Atlanta, GA, USA; Center for Machine Learning, Georgia Institute of Technology, Atlanta, GA, USA; Department of Neuroscience, Northwestern University, Chicago, IL, USA; Department of Biomedical Engineering, Northwestern University, Evanston, IL, USA; Department of Physical Medicine and Rehabilitation, Northwestern University, Chicago, IL, USA; Shirley Ryan AbilityLab, Chicago, IL, USA; Department of Neurosurgery, Emory University, Atlanta, GA, USA

## Abstract

Intracortical brain-computer interfaces (iBCIs) restore motor function to people with paralysis by translating brain activity into control signals for external devices. In current iBCIs, instabilities at the neural interface result in a degradation of decoding performance, which necessitates frequent supervised recalibration using new labeled data. One potential solution is to use the latent manifold structure that underlies neural population activity to facilitate a stable mapping between brain activity and behavior. Recent efforts using unsupervised approaches have improved iBCI stability using this principle; however, existing methods treat each time step as an independent sample and do not account for latent dynamics. Dynamics have been used to enable high performance prediction of movement intention, and may also help improve stabilization. Here, we present a platform for Nonlinear Manifold Alignment with Dynamics (NoMAD), which stabilizes iBCI decoding using recurrent neural network models of dynamics. NoMAD uses unsupervised distribution alignment to update the mapping of nonstationary neural data to a consistent set of neural dynamics, thereby providing stable input to the iBCI decoder. In applications to data from monkey motor cortex collected during motor tasks, NoMAD enables accurate behavioral decoding with unparalleled stability over weeks-to months-long timescales without any supervised recalibration.

## Introduction

In people with paralysis, intracortical brain–computer interfaces (iBCIs) provide a pathway to restoring voluntary movements by interfacing directly with the brain to translate movement intention into action^1,2^. iBCIs use implanted electrodes to record activity from populations of neurons and decoding algorithms to translate the recorded activity into control signals for external devices. In recent years, iBCIs have attained impressive performance in a range of applications, including the control of anthropomorphic robotic arms, stimulation of paralyzed muscles to enable reaching and grasping, and even rapid decoding of handwriting^3–6^.

Despite these impressive demonstrations, a key challenge limiting the clinical deployment of iBCIs is their robustness to neuronal recording instabilities that cause changes in the particular neurons being monitored over time^7–10^. Recording instabilities are attributed to a variety of phenomena, including shifts in electrode positions relative to the surrounding tissue, electrode malfunction, cell death, and physiological responses to foreign materials. As the particular neuronal population being monitored changes, so does the relationship between recorded neural signals and intention, which creates a non-stationary input to the iBCI’s decoder. Without appropriate compensation, iBCI use must be periodically interrupted to perform supervised decoder recalibration, in which neural data are collected while subjects attempt pre-specified movements. This process can be required once or even multiple times per day to maintain high-performance^2^, obstructing activities of daily living and creating additional burdens for iBCI users. Because the reliability of assistive devices is a key predictor of real-world use^11^, iBCI instabilities are often cited as motivation for alternate neural interfaces such as electrocorticography, which offer more limited but potentially more stable performance^12^.

Automatic, unsupervised decoder recalibration would provide a means to compensate for neural interface instabilities using only neural data collected during normal iBCI device use, thus preserving performance without interrupting use. One promising avenue to unsupervised recalibration is the use of latent, network-level properties of neural activity^13–16^. In particular, a few recently-developed iBCIs leverage latent manifolds, revealed by the patterns of co-activation within the neuronal population, as the foundation for a more stable neural interface^9,17–19^. Manifold-based iBCI decoders use a two-stage approach: first, a *neurons-to-manifold* mapping that transforms recorded neuronal population activity onto the underlying manifold and second, a *manifold-to-behavior* mapping that transforms manifold activity into intended movements^17–20^. Because manifolds are independent of the specific neurons being recorded, different sets of recorded neurons can be mapped onto the same manifold^17–22^. And because these manifolds have a consistent relationship with behavior extending even to years^20,22^, stable decoding can be achieved by properly recalibrating the neurons-to-manifold mapping without changing the manifold-to-behavior mapping.

A complementary avenue to improve the performance and stability of iBCI decoders is to incorporate latent dynamics, or the rules that govern the evolution of population activity over time^23^. Models of neural population activity that incorporate dynamics have already shown promise for improving iBCI performance, as they produce representations that are informative of behavior on a moment-to-moment basis and millisecond timescale^21,24–27^. Dynamics may also be useful for improving stability because dynamics, like manifolds, have a stable relationship with behavior for months to years and are independent of the specific population of neurons being monitored within a given area^21,22^. To date, however, unsupervised efforts to stabilize iBCI decoding have not incorporated this temporal information.

Here we test Nonlinear Manifold Alignment with Dynamics (NoMAD), a novel platform for unsupervised stabilization of iBCI decoding. NoMAD uses a manifold-based iBCI decoder that incorporates a recurrent neural network model of dynamics. As instabilities cause changes in the recorded neural population, the learned dynamics model can be used to help update the neurons-to-manifold mapping without knowledge of the subject’s behavior.

We applied NoMAD to recordings from monkey primary motor cortex (M1) collected during motor tasks in sessions that span multiple weeks and compared it to two previous state-of-the-art stabilization approaches that use latent manifolds. When applied to recordings from a monkey performing a two-dimensional isometric wrist force task, NoMAD achieved strikingly higher decoding performance compared to previous approaches without noticeable degradation over 3 months. Further, when applied to recordings from a behavior with very different output dynamics—a center-out reaching task, with sessions spanning 5 weeks—NoMAD again achieved substantially higher decoding performance and stability than previous approaches. These results demonstrate that unsupervised decoder recalibration using dynamics can greatly extend the time spans over which stable iBCI use is feasible and provides a new pathway to more practical iBCIs.

## Results

### Leveraging manifolds and dynamics to stabilize iBCI decoding

We begin with a conceptual schematic (**Fig. 1**). As with previous manifold decoding approaches^17–19,28,29^, our approach starts with a supervised training dataset containing neural activity and movement information from an initial recording session, which we call Day 0. We can use this dataset to characterize the manifold and dynamics, while also training a decoder to map manifold activity onto behavior. In some later recording period, termed Day K, neural recording instabilities have changed the specific neurons that are monitored, so electrode channels may now have a different relationship to the underlying manifold and dynamics. Thus, the original decoding axis no longer reflects the relationship between the manifold and behavior. As with other unsupervised methods, the high-level goal of NoMAD is to compensate for recording instabilities by learning a mapping from the Day K data onto the original manifold, allowing the original Day 0 decoder to be used. Unlike previous methods, NoMAD uses information about the temporal evolution of neural activity to help learn this mapping.

**Fig. 1.**
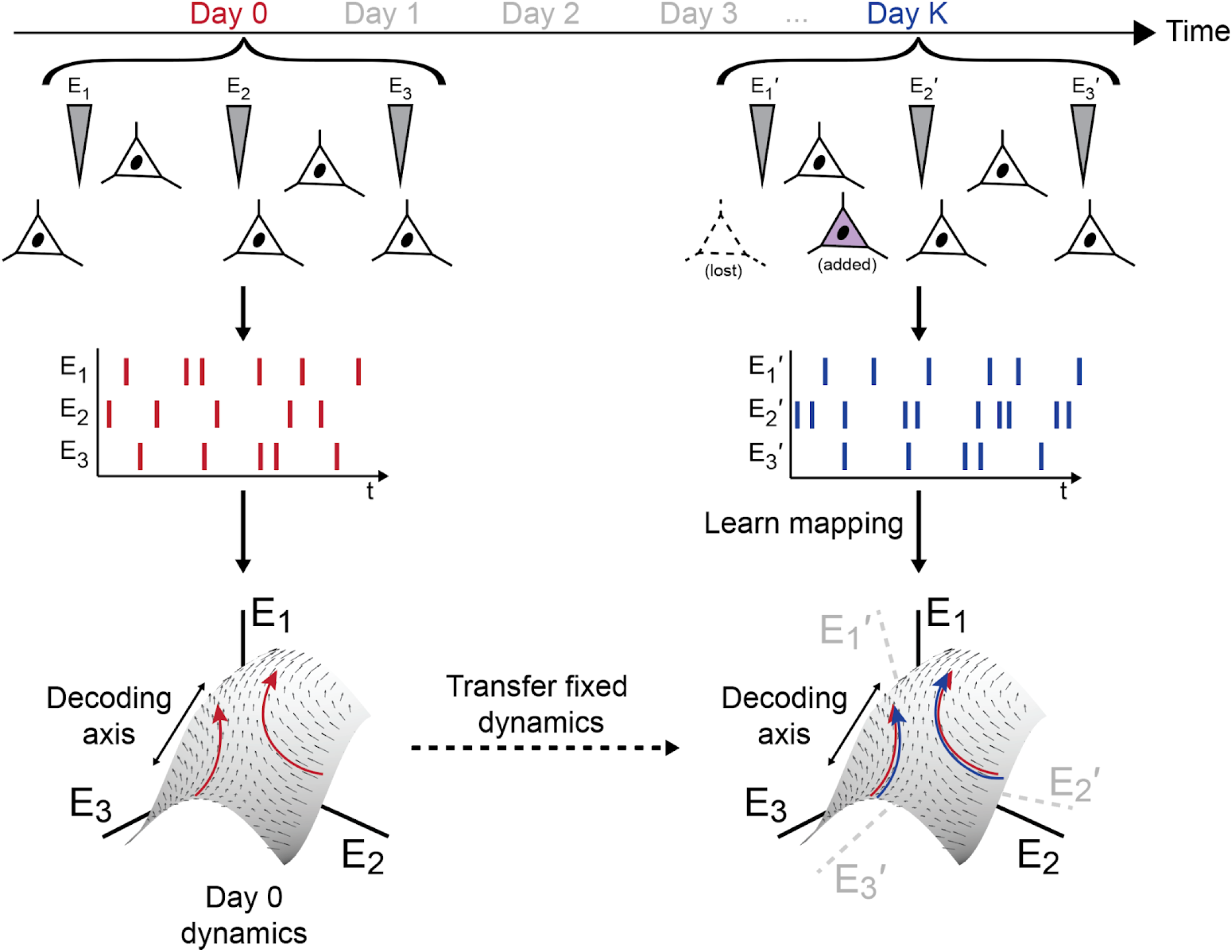
Manifold alignment with dynamics can stabilize representations of neural activity despite changes in recording conditions. We begin with a supervised training dataset containing neural activity and corresponding behavior from an initial recording session (*Day 0*). For a 3-electrode example (electrodes E_1_, E_2_, E_3_), population activity exhibits underlying manifold structure in a 3-D neural state space in which each axis corresponds to the firing rate from a given electrode. The evolution of population activity in time exhibits consistent dynamics (vector field). The relationship between manifold activity and behavior, for simple linear decoding of a hypothetical 1-D behavioral variable, is represented by a *Decoding axis*, which is assumed to be consistent over time. In a subsequent recording session (*Day K*), instabilities lead to changes in the recorded neural population, and the Day K activity (E_1_’, E_2_’, E_3_’) has a different relationship to the underlying manifold, dynamics, and decoding axis (schematized by a rotation). With NoMAD, our goal is to learn a mapping from the Day K neural activity to the original manifold and dynamics in an unsupervised manner. This allows the original decoding axis to be applied to accurately decode behavior.

### Adapting the LFADS architecture for manifold alignment using NoMAD

NoMAD models dynamics using latent factor analysis via dynamical systems (LFADS; **Fig. 2a**), a modification of standard sequential variational autoencoders (VAEs)^30–32^ which has been previously detailed^21,30,33^. Briefly, LFADS approximates the dynamical system underlying an observed neural population using a recurrent neural network (RNN; the “Generator”) that receives a sequence of inferred inputs. Additional RNNs encode the initial state of the dynamical system and infer the sequence of inputs to the Generator. The model outputs firing rate predictions for the observed neurons through a linear readout from the Generator—this readout sets the relationship between the learned manifold and the high-dimensional neural activity (i.e., it orients the manifold within the high-dimensional neural state space). In standard LFADS, the model training objective is to maximize a lower bound on the marginal likelihood of the observed spiking activity, given the firing rates output by the model. All weights in the model are updated during training via backpropagation through time.

**Fig. 2.**
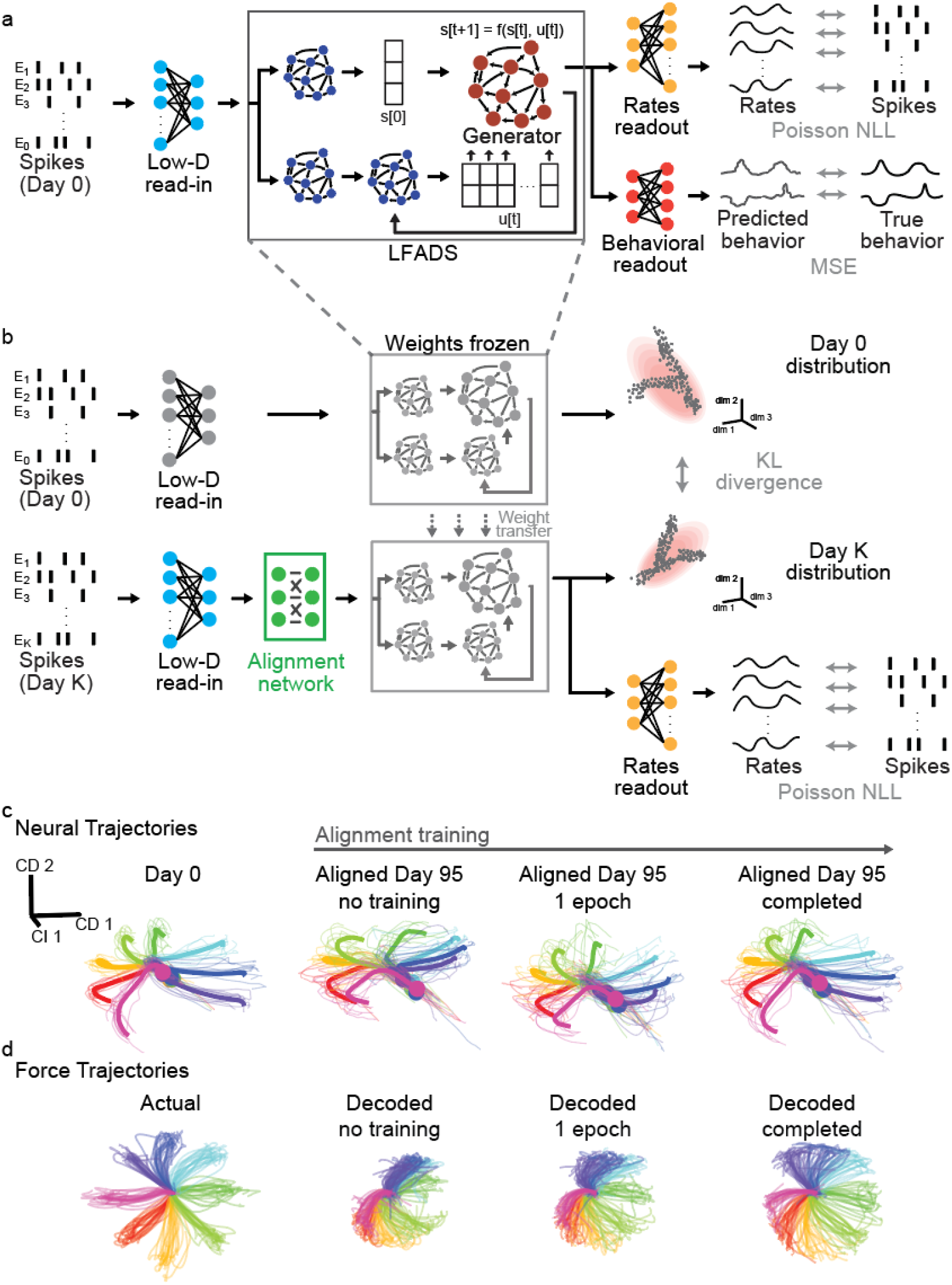
Modeling and aligning the neural manifold using NoMAD. (a) We first train an LFADS model on a given reference Day 0 to estimate the manifold and dynamics from recorded spiking activity. LFADS models neural population dynamics using a series of interconnected recurrent neural networks (RNNs). All of the model parameters are trained by simultaneously minimizing the reconstruction losses of the spiking activity (Poisson negative log-likelihood) and the recorded behavior (mean squared error). (b) To apply the same LFADS model on a later Day K, we freeze the parameters of the trained LFADS model, and introduce a feedforward “Alignment network” to transform the new day’s spiking activity to be compatible with the previously trained LFADS model. The Alignment network is trained by simultaneously minimizing the KL divergence between the distributions of the Day 0 and Day K Generator states and the reconstruction loss of the spiking activity (Poisson NLL). (c) Dimensionality reduction applied to the Generator states for Day 0, and over the course of alignment training for data collected 95 days later. Neural data were recorded from M1 as a monkey performed an isometric force task. Colors indicate trajectories for the different target locations. Thick lines indicate the trial average, and thin lines denote single trials (10 representative trials per condition are shown here). Initial estimates of Day 95 Generator states converged over training to resemble Day 0 Generator states, allowing the same decoder to achieve high accuracy force predictions. (d) Left: Measured single trial force trajectories, colored by target location. Right: decoded forces across several epochs of Day 95 alignment to Day 0: before alignment (R^2^ = 0.64), after 1 epoch of alignment training (R^2^ = 0.73), and after completed alignment training (116 epochs; R^2^ = 0.86).

To model the Day 0 supervised training dataset, we made two main modifications to the LFADS architecture. First, we added a low-dimensional read-in matrix at the model’s input to standardize the dimensionality of the input to the RNNs. This read-in allows the same LFADS model to be applied to Day K datasets despite changes in the number of recorded electrodes from Day 0, such as those which may occur due to the removal of channels with sparse activity by the experimenter. Second, we added a readout matrix that predicts behavioral data from the Generator’s activity, similar to methods that aim to learn dynamics that are related to both the recorded neural activity and measured behavioral variables^34^. Behavioral prediction serves as a second training objective, adding a source of information that is complementary to the observed spiking activity, which helps ensure that the model converges to a good solution^34,35^. After training the LFADS model, we learn the manifold-to-behavior mapping, i.e., a decoder that predicts the Day 0 behavioral data from the Day 0 Generator states. Separating training of the LFADS model and Day 0 decoder allows the use of decoding architectures with more complexity and capacity than a simple, single-time-step linear decoder (e.g., Wiener filters, RNNs, etc.) without impacting the learned manifold or dynamics.

Because neural dynamics and behavior have a stable relationship over months to years^21,22^, both the Day 0 neural dynamics model and decoder should be applicable to data collected after the initial supervised dataset. However, because recording instabilities distort the relationship between recorded neurons and the manifold, we must periodically update the mapping between the neural activity and the manifold through an alignment transformation, which fortunately, can be an unsupervised process. Once this transformation is learned, data on a subsequent Day K can be passed through the Day 0 dynamics model, such that the Day 0 decoder can be used to predict Day K behavior with high accuracy.

To update the Day K neurons-to-manifold mapping in NoMAD, the weights of the LFADS RNNs, including the Generator that expresses the latent dynamics, are held constant, while three other network components are learned or updated with an unsupervised alignment step: a feedforward alignment network that adjusts the input to the RNNs, the low-D read-in, and the rates readout (**Fig. 2b**). During the alignment step, there are two training objectives: 1) minimizing the difference between the distributions of the Generator states on Day 0 and Day K, and 2) maximizing the likelihood of observed Day K spiking activity given the firing rates output by the model. At each training step, the Day 0 and Day K Generator state distributions are compared by first approximating each by a multivariate normal distribution and then computing the Kullback-Leibler (KL) divergence between those normal distributions. The multivariate normal approximation focuses the alignment process on matching first and second order statistical moments of the Generator state distributions, making the alignment problem more tractable than matching higher order statistics. Model weights that are adjusted during alignment are updated via backpropagation through time.

To illustrate how NoMAD adjusts the manifold and dynamics, we applied it to twenty datasets collected from a monkey performing an isometric force task over the course of 95 days. In this task, neural data were recorded from a 96-channel electrode array in M1 while a monkey generated wrist forces to control an on-screen cursor, with the goal of reaching a target in one of eight directions on each trial. We visualized the low-D structure of the Generator states by finding a 3-dimensional subspace via demixed principal component analysis (dPCA)^36^. We first fit dPCA parameters to the Generator states inferred by the LFADS model from Day 0 neural activity. We then applied those dPCA parameters to the Generator states on Day K across three phases of unsupervised NoMAD alignment: before any training had occurred, after one training epoch, and after the completion of training (**Fig. 2c**). By using the same dPCA parameters on Day 0 and Day K, we could directly compare the low-D trajectories across Day 0 and Day K. dPCA revealed a structure underlying Day 0 activity that clearly distinguished neural activity from the 8 different target conditions. Prior to alignment, the low-D structure of the Generator states on Day K was distinct from the structure on Day 0, as expected. After alignment with NoMAD, the low-D structure on Day K more closely matched Day 0. We then tested whether this unsupervised alignment led to greater decoding stability by first training a Wiener filter decoder that mapped the Generator states to the measured forces on Day 0, and then testing this decoder on Day K data over several epochs of NoMAD alignment. As shown, the alignment procedure improved force decoding substantially (**Fig. 2d**).

### NoMAD stabilizes offline decoding during an isometric force task

We applied NoMAD to pairs of recording sessions from a monkey performing the isometric force task over 95 days (**Fig. 3a**). Behavioral decoding consisted of a Wiener filter that mapped manifold activity onto the recorded 2-D forces with high temporal resolution (20 ms time steps). The Day 0 dynamics model (LFADS) and Day 0 decoder (Wiener filter) were trained on a supervised dataset from a single session, and we evaluated NoMAD’s ability to enable accurate and stable decoding performance on a different session (Day K) through unsupervised alignment. We compared NoMAD against a standard Wiener filter decoder that was trained using smoothed spiking activity and behavior from Day 0 and evaluated on Day K without any adjustment (*Static decoder*), and also to two state-of-the-art manifold-based stabilization techniques: aligned factor analysis (Aligned FA)^17^, which is based on linear dimensionality reduction, and the adversarial domain adaptation network (ADAN)^18,19^, which uses a neural network autoencoder for dimensionality reduction and generative adversarial networks for alignment. We note that NoMAD alignment is completely unsupervised, in that we use all available data from a session, including periods where the monkey was inactive. For Aligned FA and ADAN, we use only within-trial data where the monkey is active, to be consistent with the original demonstrations of these methods. We quantified Day K force decoding using the coefficient of determination (R^2^), i.e., the fraction of variance of the recorded force signal on Day K that is predicted by the decoder. We term evaluations with negative R^2^ to be decoding failures.

**Fig. 3.**
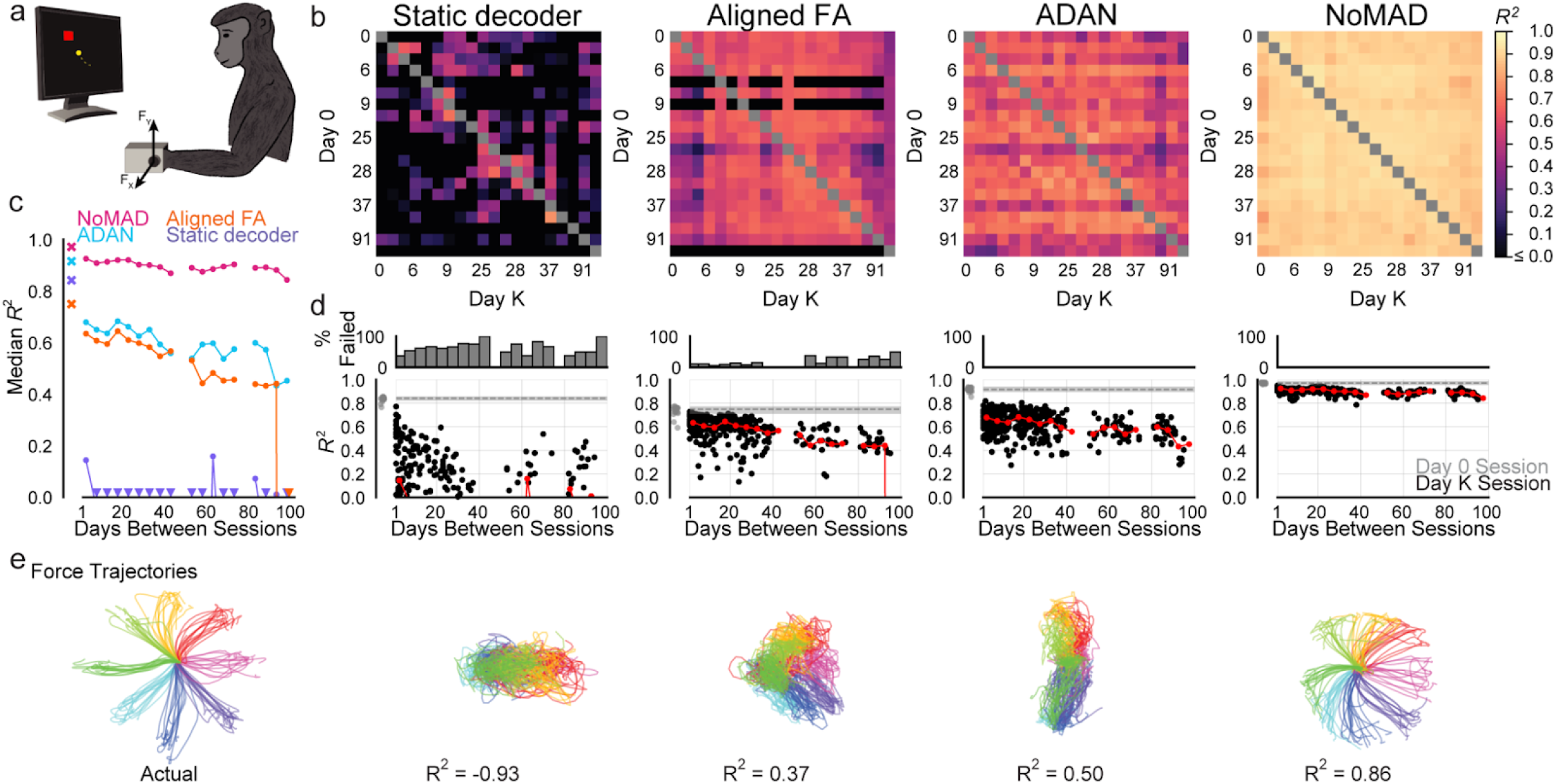
NoMAD enables stable offline decoding across months in an isometric force task. (a) Schematic of isometric force task. (b) R^2^ of aligned force decoding for each method applied to all pairs of sessions (20 sessions spanning 95 days). (c) Median R^2^ of force decoding within each 5-day bin. Triangles indicate points that fall below the limit of the y-axis and x-markers indicate median within-day force decoding performance on Day 0. (d) Top: Percentage of Day K decoding failures (R^2^<0) in 5-day bins. Bottom: Decoding performance as a function of days between sessions. Black points indicate Day K decoding performance for single pairs of days. Red points show the median Day K performance within each bin of width 5 days. Gray points denote initial performance for Day 0 decoders (i.e., evaluated on held-out same-day data) for each of the 20 datasets. Gray dashed lines indicate median decoding performance of Day 0 decoders, and shaded regions around them represent the first and third quartiles. (e) Left: Measured single trial force trajectories on Day 95, colored by target location. Right: decoded forces for all methods when Day 95 is aligned to Day 0.

Prior to evaluating across-session decoding stability, we first evaluated baseline decoding performance for each method by training Day 0 decoders for each session and testing those decoders on held-out data from the same session. Day 0 decoders used for NoMAD, which were trained on Day 0 LFADS generator states, achieved the highest median performance and had the least amount of variability for the twenty sessions (median R^2^ [Q1, Q3] = 0.972 [0.967, 0.976]), followed by ADAN (0.916 [0.908, 0.929]). Aligned FA (0.749 [0.722, 0.761]) and the Static decoders (0.842 [0.834, 0.846]) had lower and more variable within-day decoding performance. These initial Day 0 performance metrics serve as a useful upper-bound in interpreting the efficacy of each alignment method.

To measure decoding stability across a wide variety of recording conditions, we tested each method on all pairs of sessions (380 pairs; **Fig. 3b**). As expected, static decoders trained on a given session failed to generalize to other sessions. This resulted in rapid degradation of decoding performance over time and frequent decoding failures, such that the median decoding performance was quite poor for pairs of sessions spaced less than 5 days apart (R^2^ = 0.14) and negative for pairs spaced further apart (223 decoding failures; **Fig. 3c, d**).

Aligned FA achieved more stable decoding than the static decoder across time (half life = 45.1 days), but with only moderate performance (median R^2^ = 0.59) and high variability. In addition, Aligned FA repeatedly failed for most alignments that were initialized using particular sessions (shown by the black horizontal bars in the heatmap), resulting in 51 total decoding failures. Separately, we also attempted to apply Aligned FA in a sequential fashion, i.e., performing alignment between sequential recording sessions to try to maintain stability (the approach described in the original study), but found that this further decreased performance and increased variability (results not shown). ADAN achieved greater performance improvements (median R^2^ = 0.65) and stability (half life = 76.7 days), with 0 decoding failures. Yet, performance for individual pairs of sessions was highly variable.

Compared to these previous methods, NoMAD achieved strikingly higher performance (median R^2^ = 0.91) with little variability, no failures, and only modest performance degradation across the 3-month window (half life = 209.1 days). These differences were also evident when visualizing the decoded data: single-trial forces decoded by NoMAD were more consistent with the measured forces than those produced with other methods (**Fig. 3e**).

### NoMAD stabilizes offline decoding during an unloaded reaching task

To ensure that NoMAD’s efficacy extends beyond the isometric task, we also evaluated its performance on recording sessions from a monkey performing a center-out reaching task^20,37^. On each trial, a monkey used a manipulandum to move a cursor from the center of a screen to one of eight targets spaced equally around a ring (**Fig. 4a**). We applied NoMAD to 96 channels of data recorded from M1 over 12 sessions spanning 38 days. As in the isometric task, we used a supervised dataset to train the Day 0 LFADS model and Wiener filter decoder. We evaluated NoMAD on a separate Day K dataset by performing unsupervised alignment prior to applying the Day 0 decoder. We again tested NoMAD’s performance against ADAN, Aligned FA, and a Static decoder on the same data^17–19^. Day K decoding was quantified using the R^2^ between the recorded and predicted cursor velocity signals, and negative R^2^ values were again classified as decoding failures.

**Fig. 4.**
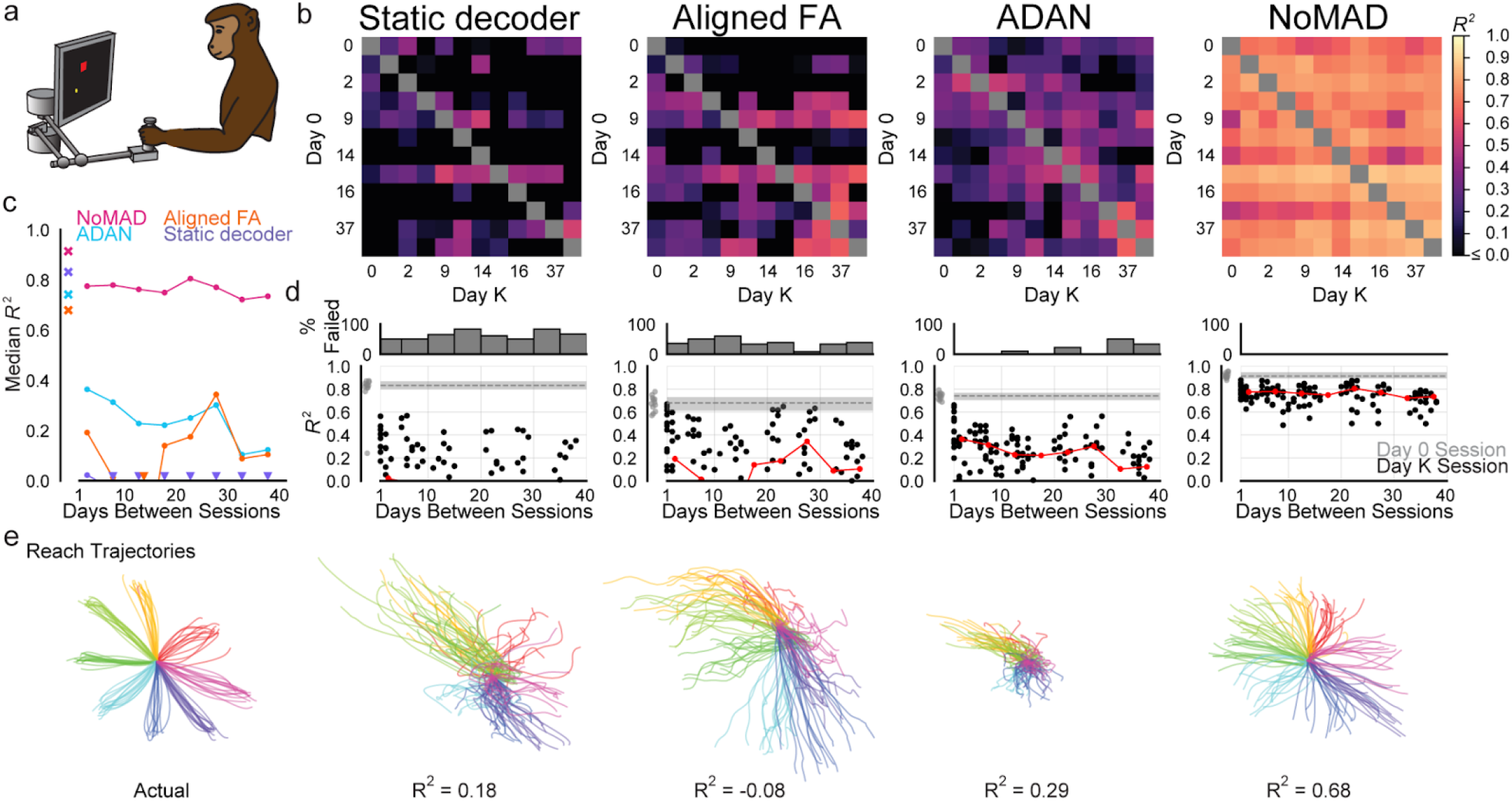
NoMAD enables stable offline decoding across 5 weeks in a reaching task. (a) Schematic of the center-out reaching task. (b) R^2^ of aligned cursor velocity decoding for each method applied to all pairs of sessions (12 sessions spanning 38 days). (c) Median R^2^ of cursor velocity within each 5-day bin. Conventions as in Fig. 3. (d) Top: Percentage of Day K decoding failures (R^2^<0) in 5-day bins. Bottom: Decoding performance as a function of days between sessions. Conventions as in Fig. 3. (e) Left: Measured single trial reach trajectories, colored by target location. Right: reach position trajectories integrated from the decoded reach velocity when Day 38 is aligned to Day 0.

We again started by evaluating baseline decoding performance for all sessions using decoders trained on Day 0 data and evaluated on held-out Day 0 data for each method. NoMAD achieved the highest and least variable performance amongst the twelve sessions (median R^2^ [Q1, Q3] = 0.914 [0.906, 0.936]). Static decoders fit to smoothed spikes were the next highest performing for this dataset, but were highly variable (0.832 [0.813, 0.856]). ADAN within-day decoders’ performance was lower but had variability similar to NoMAD (0.742 [0.716, 0.757]). Aligned FA yielded initial decoders with the lowest performance and highest variability on this task (0.679 [0.624, 0.718]).

We next assessed whether each method could facilitate stable decoding across sessions by testing alignment on all pairs of days (132 pairs; **Fig. 4b**). Static decoders again failed to yield accurate decoding across sessions, with poor median performance even for sessions separated by less than five days (R^2^ = 0.022), and primarily negative performance thereafter. A total of 78 pairs of days result in decoder failures (**Fig. 4c, d**).

This dataset was more challenging to align for Aligned FA and ADAN than was the isometric task. Aligned FA provided some accuracy improvement over the static decoder (median R^2^ = 0.17). Again, there was significant variation in decoding performance, which resulted in 53 decoding failures and a clear decline in performance (half life = 1.03 days). ADAN provided some additional improvement in terms of accuracy (median R^2^ = 0.29) and stability (half life = 7.12 days, 15 decoding failures) but still showed high variability amongst performance for individual pairs.

NoMAD exhibited higher decoding performance (median R^2^ = 0.77) with no decoding failures and less variability in performance over the 5-week timespan. NoMAD also showed less degradation in decoding accuracy (half life = 61.4 days). Visualizations of the decoded cursor trajectories confirmed that NoMAD produces more consistent cursor velocity estimates following stabilization than do other methods (**Fig. 4e**).

## Discussion

We introduced NoMAD, an unsupervised manifold alignment technique that leverages manifolds and their dynamics to achieve stable decoding over long timespans. In our tests, NoMAD improved both decoding accuracy and stability over 95 days in an isometric wrist task and over 38 days in a reaching task. Other methods resulted in less decoding improvement or marked performance degradation over time. By incorporating the temporal structure of the neural activity into the manifold-alignment process via dynamics, NoMAD enables more stable neural decoding that may require less frequent iBCI recalibration procedures.

### Relation to Previous Work

A variety of strategies have been used to reduce the reliance on supervised decoder recalibration^10,17–19,28,29,38^. One approach uses neural network decoders and months-long datasets that expose the decoder to a wide variety of recording instabilities to learn a mapping from neuronal population activity to movement intention that is robust to changes in the recorded neurons^10^. However, collecting such large supervised datasets requires a substantial time commitment from the user and is therefore challenging to perform clinically. A second strategy, “semi-supervised” recalibration, involves automatically adapting the decoder on the fly using a retrospective analysis of data collected during the subject’s normal use of the iBCI^38^. For example, in a setting with predefined targets, the neural activity preceding movement to a given target likely reflects the user’s intention to move toward that target. This data can then be used to update the decoder, much like supervised decoder training. However, this strategy only works when the user’s intent can be guessed post-hoc—as in BCI spellers or movement among a limited number of predefined targets—and is thus unlikely to scale to more complex and naturalistic settings.

Another manifold alignment approach, Distribution Alignment Decoding (DAD), is conceptually similar to the approach used here, but performs alignment of low-dimensional activity from two datasets by doing a brute force search of candidate rotations in the low-D space^28^. However, this alignment approach fails when distributions of movements are symmetric. Further, when dimensions of neural activity scale beyond extremely simple representations (e.g., beyond 2 or 3 dimensions), a brute force search quickly becomes intractable. More recently, Hierarchical Wasserstein Alignment (HiWA) improves on this approach using neural networks^29^, but relies on the existence of discrete structure, such as clusters within the neural activity that represent similar movements.

Our method improves upon previous manifold-alignment efforts by incorporating temporal constraints via nonlinear dynamics models. In addition, the method is entirely unsupervised, in contrast to methods such as canonical correlation analysis (CCA) and previous methods that exploit dynamics to improve iBCI longevity^20,22^.

A practical limitation of the approach is that the NoMAD alignment process, like most neural network training, is computationally demanding. Thus without further optimization for implanted medical devices with severe power constraints, the alignment process is best run on external computing hardware. Despite such constraints, the benefits of NoMAD to stability and accuracy indicate a promising advance to iBCIs.

### Potential Future Applications

In this work we test the NoMAD approach with particularly rigorous constraints, specifically: 1) the alignment process must be completely unsupervised (i.e., cannot incorporate any behavioral information), and 2) only a small (minutes-long) initial supervised training dataset is used to calibrate the model. Even with these requirements, we found that NoMAD could stabilize decoding performance for many weeks to months. However, these constraints could clearly be relaxed in a practical application. For example, assuming periods of reasonable decoding stability, data collected during BCI use comes with an inherent set of behavioral labels (i.e., we know how the subject was using the BCI), and this information could be incorporated to guide the alignment process. Similarly, as more and more datasets are collected during BCI use for a given subject, one could train an aggregated model that spans those datasets (similar to previous work^10,21^), which would provide decoders that become inherently more stable as more data is collected, thus lessening the need for frequent recalibration.

We note that while the current work focuses on spiking activity recorded via intracortical electrode arrays, this is not an inherent limitation of the approach. Indeed, LFADS, upon which NoMAD is based, has been applied successfully to improve behavioral state classification from electrocortographic (ECoG) recordings^39^, which suggests that the NoMAD approach could be made to generalize to other BCI recording modalities and signal sources.

Recent effort has been focused on improving the inherent stability of electrode arrays, for example through biomaterials and tissue engineering innovations that reduce the inflammatory effect on surrounding tissue, promote cell growth, and enable flexible movement with the brain^40–45^. However, with electrode arrays currently approved for chronic implantation in human subjects, clinical iBCIs remain subject to neural interface instabilities that necessitate algorithmic solutions^2^. As new neural interfaces progress to clinical applications, they are likely to complement algorithmic stabilization approaches, resulting in improved stability beyond what is possible with either approach individually.

A potential limitation of current manifold alignment methods is that they rely on a stable relationship between activity on the neural manifold and behavior over time. While this assumption is reasonable for previously-learned behaviors^20,21^, it may not be the case if subjects are learning new skills. Indeed, though learning need not change the manifold on short timescales^46^, long-term learning likely results in manifold-level changes^47^, which might also affect the relationship between manifold activity and behavior. However, as demonstrated by several methods including ours^17–19,28,29^, stabilization does not require massive data libraries (e.g., spanning many months), but instead can be achieved using recording sessions lasting only tens of minutes. Therefore, if the latent manifold and dynamics are changing due to processes like learning, stabilization could be run at shorter temporal intervals over which the manifold and dynamics should be locally stable. Further studies in this area could help determine whether and how manifold-alignment techniques need to account for learning.

Another limitation of the tests of manifold stabilization approaches to date is the consistency of behavior in the datasets used. The behavioral consistency achieved in the lab setting assured that the datasets were rich enough and similar enough to be alignable. In practical applications, a key assumption of all stabilization approaches is that iBCI decoding performance will be stable for certain time periods, such that data can be collected during use of the iBCI and periodic alignment can be performed to maintain manifold stability. If decoding performance is stable for reasonable time periods without alignment (e.g., many hours), this could ensure that the datasets cover rich enough and similar enough behavioral distributions (e.g., by spanning large enough regions of behavioral space) for successful periodic alignment.

An open neuroscientific question is the degree of similarity in manifold structure and dynamics across behaviors. This question has profound implications for building BCIs that generalize across behaviors. Recent studies suggest that different behaviors may occupy distinct manifolds^48^. As such, BCIs that rely on manifold structure may require different mappings from manifold activity to behavior, and would need to adjust their decoding depending on the manifold or manifolds that are currently occupied. This also affects stabilization strategies—to date, manifold-based stabilization methods have been tested only on datasets containing single behaviors. However, solutions to address the multiple-manifold scenario exist, including labeling data collected during BCI use by the behavior that was being performed and using that information to guide alignment.

Our offline measures of NoMAD’s performance demonstrate the potential of neural population dynamics to yield unprecedented stability and accuracy for iBCI decoders. Future investigation into its benefits as an unsupervised recalibration procedure during online iBCI use will further illuminate how it may impact device development and lead to more feasible real-world devices.

## Acknowledgements

We thank Matthew Perich and Stephanie Naufel for the collection of the data used in this work. We thank Ali Farshchian for helpful discussions regarding ADAN and Fabio Rizzoglio for helpful discussions regarding ADAN and Aligned FA. This work was supported by the Emory Neuromodulation and Technology Innovation Center (ENTICe), NSF NCS 1835364, DARPA PA-18-02-04-INI-FP-021, NIH Eunice Kennedy Shriver NICHD K12HD073945, NIH-NINDS/OD DP2NS127291, NIH BRAIN/NIDA RF1 DA055667, the Alfred P. Sloan Foundation, the Burroughs Wellcome Fund, and the Simons Foundation as part of the Simons-Emory International Consortium on Motor Control (CP), NIH NINDS NS053603 and NS074044 (LEM), NIH NIBIB T32EB025816 (BMK, YHA), NSF Graduate Research Fellowship DGE-2039655 (ARS).

## Code availability

Code will be made available upon publication.

## Data availability

Data will be made available upon publication.

## Author Contributions

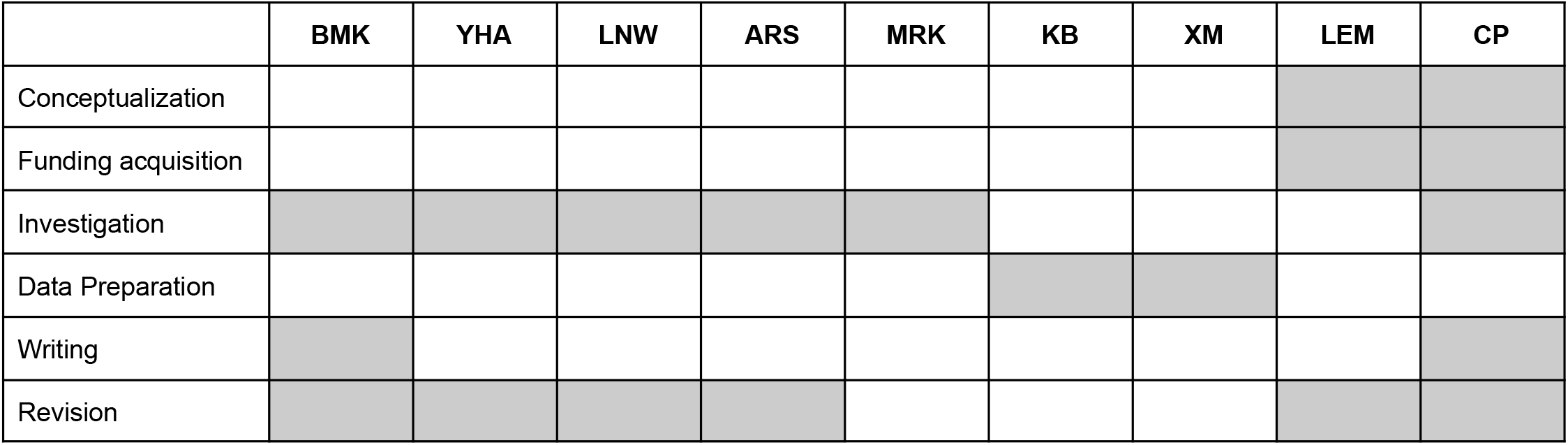

## Methods

### Monkey Surgical Implants

After training animals to perform the tasks, a 96-channel microelectrode array with 1.5mm-long electrode shanks (Blackrock Microsystems, Salt Lake City, Utah) was implanted into the hand area of primary motor cortex (M1). Prior to implanting the array, the hand area of M1 was identified intraoperatively through sulcal landmarks and by stimulating the surface of the cortex to elicit twitches of the wrist and hand muscles.

### Monkey Data Collection

#### Collection

Cortical data was collected using the Cerebus data acquisition system from Blackrock Microsystems. Raw cortical data was collected at 30kHz and filtered with a 250Hz high-pass filter. Timestamps of when the filtered data passed below a provided threshold were recorded as threshold crossings, commonly also called spikes. This process was all done using software on the Cerebus system.

Force data for the Isometric Task was collected at 2kHz using the Cerebus analog inputs. Manipulandum joint data for the Reaching Task was collected using a NI DAQ card connected to a Mathworks XPC system that controlled the task. The XPC system calculated the endpoint coordinates of manipulandum, then sent the coordinates to the Cerebus through a series of digital packets.

#### Binning

The data was binned into 1ms segments. For the threshold crossings, the number of spikes per bin was counted. For the force and manipulandum position data, the data was filtered and downsampled to 1kHz. The cortical and task bins were aligned in time.

### Monkey Isometric Task

Monkey J was trained to operate a 2D isometric wrist force device. The monkey’s left arm was positioned in a splint to immobilize the forearm in an orientation midway between supination and pronation (with the thumb upwards). A small box was placed around the monkey’s open left hand, incorporating a six-axis load cell aligned with the wrist joint. The box was padded to comfortably constrain the monkey’s hand and minimize its movement within the box. The monkey controlled the position of a cursor displayed on a monitor by the force exerted on the box. Flexion and extension force moved the cursor right and left, respectively, while forces along the radial and ulnar deviation axis moved the cursor up and down. Prior to placing the monkey’s hand in the box, the force was nulled in order to place the cursor in the center target. Targets were displayed either at the center of the screen (zero force), or equally-spaced along a ring around the center target.

The monkey performed a center-out task that began with the appearance of the center target. The monkey was allowed two seconds to move to the center target and was required to hold for a time randomly chosen from a uniform distribution between 0.2s and 1.0s. A successful center hold triggered the appearance of one of eight possible outer targets, chosen in a block-randomized fashion. The monkey was allowed another two seconds after target onset to move the cursor to the outer target. The required hold time for the outer target was 0.8s. Successful trials ended with the delivery of a liquid reward. Failure to reach a target within the allowed two seconds or to remain within a target as required, resulted in an aborted (center target) or failed (outer target) trial. Successive trials were separated by a two-second inter-trial interval. This data has been published in Ma*, Rizzoglio*, et al., *bioRxiv*.

### Monkey Reaching Task

Monkey C was trained to make reaching movements using a planar manipulandum in a two-dimensional center-out manner. To begin each trial, the monkey moved his hand to the center of the workspace. After a waiting period, the monkey was presented with one of eight equally spaced outer targets, arranged in a circle and selected uniformly at random. The monkey was trained to hold for a variable delay period, after which he received an auditory go cue. After the go cue, the monkey had 1s to reach the outer target and hold within the target for 0.5s. A successful trial led to a liquid reward. The cursor position was recorded at 1kHz using joint encoders. Timing events such as go cues were logged digitally. This data was previously published in Perich et al., *Neuron* 2018; Gallego, et al., *Nature Neuroscience* 2020; and Ma*, Rizzoglio*, et al., *bioRxiv*.

### Data Preprocessing

We apply the following preprocessing steps to the data for all methods compared:

#### Resampling

In order to resample the data to larger bin sizes than originally provided (e.g. to work with 20ms bins from 1ms bins), we must handle both spiking data and continuous-valued data. For spiking data, we aggregate bins by summing the number of spikes. For continuous-valued data, we apply a Chebychev filter (order = 500) for anti-aliasing, and then downsample the data to the appropriate sampling rate.

#### Highly Correlated Channel Removal

In order to prevent overfitting to correlated noise events across channels, we remove channels that are involved in many high correlations with other channels. We first compute the cross correlations between all pairs of channels. We set a threshold above which we want to remove correlations. For the monkey datasets, we set a threshold of 0.2 (computed when the data is in 1ms bins). We remove any channel that is involved in a correlated pair above this threshold.

#### Behavioral Outlier Removal

For datasets with continuous behavioral variables (e.g. force, cursor position), behavior was evaluated for values that fall far outside of the distribution of values for a typical trial. These outlier values can lead to errors in training the behavioral readout of the LFADS models, ADAN and Aligned FA models, and decoders. We determined cutoffs for removing values that were applied to all datasets during dataset loading. Force values are in units of voltage directly from the load cell, and cursor position values are in terms of the handle position in units of distance. These cutoffs are shown in Table 1.

**Table 1.** Behavioral cutoffs for outlier removal.

| Monkey | Behavioral Variable | Lower Bound, X | Upper Bound, X | Lower Bound, Y | Upper Bound, Y |
| --- | --- | --- | --- | --- | --- |
| Isometric | Force (mV) | -1500 | 800 | -1100 | 600 |
| Isometric | dF/dt (mV/t) | -100 | 100 | -100 | 100 |
| Reaching | Cursor position (cm) | 0.5 | 1.5 | -1.1 | -0.25 |
| Reaching | Cursor velocity (cm/t) | -0.025 | 0.025 | -0.025 | 0.025 |

The following preprocessing step was incorporated into the NoMAD architecture only:

#### Normalization

To account for large changes in the firing rates of individual channels across days, we normalized each channel to have zero mean and unit standard deviation. On the continuous spiking data, we first smooth the data with a 20ms Gaussian kernel. On the smoothed spiking data, we compute a per-channel mean and standard deviation. These means and standard deviations are saved for every day so that they can be applied to the LFADS input spiking data (not smoothed) before the low dimensional read-in layer. Non-normalized input data is maintained so that reconstruction cost can be computed on binned spikes.

### NoMAD Architecture and Parameters

The NoMAD architecture consists of two stages: fitting an initial core Day 0 LFADS model on training data from an initial day and aligning that model to data from a subsequent Day K.

#### Day 0 Model Architecture

The LFADS model has been detailed previously^21,30,33^. Briefly, LFADS is an instantiation of a variational autoencoder (VAE) extended to sequences. An encoder RNN (implemented using GRU units) takes as input a data sequence X(t), and produces as output a conditional distribution over a latent code Z, Q(Z|X(t)). In the VAE framework, an uninformative prior P(Z) on this latent code serves as a regularizer, and divergence from the prior is discouraged via a training penalty that scales with KL(Q(Z|X(t)) || P(Z)). A data sample is then drawn from Q(Z|X(t)), which sets the initial state of a decoder RNN. This RNN attempts to create a reconstruction R(t) of the original data via a low-dimensional set of factors F(t). Specifically, the data X(t) are assumed to be samples from an inhomogeneous Poisson process with underlying rates R(t). This basic sequential autoencoder is appropriate for neural data that is well-modeled as an autonomous dynamical system. In all applications listed, we used the modified sequential autoencoder that was adapted for modeling input-driven dynamical systems. This model contains an additional controller RNN, which compares an encoding of the observed data with the output of the decoder RNN, and attempts to inject a time-varying input U(t) into the decoder to account for data that cannot be modeled by the decoder’s autonomous dynamics alone.

The LFADS objective function is defined as the log likelihood of the data (given the Poisson process assumption above), marginalized over all latent variables. This is optimized in the VAE setting by maximizing a variational lower bound on the marginal data log-likelihood. In training, the objective function is optimized using stochastic gradient descent where the network parameters are updated through backpropagation through time. We used the Adam optimizer to optimize the objective function, and implemented gradient clipping to prevent potential exploding gradient issues. To prevent the potentially problematic large values in RNN state variables and achieve more stable training, we also limited the range of the values in GRU hidden state by clipping values greater than 5 and lower than −5.

In addition, a behavioral readout matrix is trained to learn a mapping from the generator RNN states to the continuous behavior. This matrix is trained to minimize the mean squared error between the true behavior and the predicted behavior. This objective function is again trained using stochastic gradient descent and backpropagation through time. A second Adam optimizer with its own learning rate and decay schedule is used to train this matrix.

#### Day 0 Model Training & Hyperparameters

All experimental data is modeled without regard to trial structure, i.e. the optimization process is completely unsupervised at all stages. To do so, the continuous data (an entire session) is divided into segments. We used segments of length 600 ms with 120 ms of overlap between segments. For model validation, 20% of these segments are reserved. The start indices of each trial are stored so that after training, model output can be reassembled. In order to avoid artifacts while reassembling the overlapping segments, the overlapping region is linearly scaled from 1 to 0 and added to the corresponding region in the reconstructed data, which is inversely linearly scaled (0 to 1). Non-overlapping portions of the segments are concatenated to the array normally.

In the Day 0 architecture, a few critical hyperparameters define the model, which we list in Table 2.

**Table 2.** LFADS Day 0 architecture parameters.

| parameter | Value |
| --- | --- |
| Low-D Read-In Output Dimensionality | 50 |
| Encoder RNN Units | 100 |
| Generator RNN Units | 100 |
| Controller RNN Units | 100 |
| Controller Output Dimensionality | 4 |
| Factors Dimensionality | 30 |

Datasets were fit using single LFADS models with additional behavioral readout matrices. The goal of the behavioral readout matrices was to ensure that manifolds learned on Day 0 would be highly predictive of behavior. To do this, we trained a matrix transformation from the generator states to the continuous-valued behavior. For the isometric task, the behavior being predicted by the readout included both force and the derivative of force; for the reaching task, the behavior included cursor position and velocity.

For monkey dataset experiments, single LFADS models were trained on 1 GPU each. Training stopped when there was no improvement in the performance for 10 subsequent epochs or when the learning rate reached a value of 1e-6 during the annealing process. We found hyperparameters that were able to train models for both the isometric and kinematic datasets using grid searches. We performed a grid search over a set of values for a single hyperparameter for a subset of isometric and kinematic monkey datasets to select the best value and repeated for all hyperparameters. We then trained all Day 0 models for monkey datasets using the hyperparameters in Table 3.

**Table 3.** LFADS Day 0 model hyperparameters.

| hyperparameter | value |
| --- | --- |
| Dropout rate | 0.05 |
| Coordinated dropout rate | 0.3 |
| Batch size | 1000 |
| Initial learning rate | $1e-3$ |
| Learning rate decay | 0.95 |
| L2 scale | $1e-6$ |
| L2 ramping epochs | 100 |
| KL scale | $1e-4$ |
| KL ramping epochs | 100 |
| Behavioral readout learning rate | $1e-3$ |
| Behavioral readout cost scale | 0.1 |
| Behavioral readout cost ramping epochs | 100 |

#### Data Augmentation

In order to prevent models from overfitting to individual spikes or fast oscillations in the data, a data augmentation strategy for discrete data known as ‘spike jittering’ is applied during training. In this approach, the training procedure shifts spikes randomly in time, up to 2 bins (a settable hyperparameter) before or after their original time bin, prior to modeling the data.

#### Alignment Model Training & Hyperparameters

Since we must compare data distributions from the first day to distributions from subsequent days when performing alignment, the NoMAD computational graph must contain two data flow pathways. The first pathway sends data from the first day directly through the core LFADS model. The second pathway sends data from a subsequent day through an aligner and then through the same LFADS model as the first day.

After training the LFADS model for Day 0, we trained the linear read-in matrix, the alignment network (2-layer Dense network with ReLU activations and identity initialization), the linear readout matrix from generator states to factors, and the linear readout matrix from LFADS factors to inferred firing rates for subsequent sessions. All remaining model components had weights that were held fixed.

For the base Day 0 LFADS model (without the alignment network), and Day K model (which includes the alignment network), we separately obtained the distribution of the samples from each dimension of the factors for all time points, and across the entire batch of data. We then calculated the Kullback-Leibler Divergence (KL cost) between these two full-dimensional distributions assuming they follow Multivariate Normal (Gaussian) distributions with potential correlations between each dimension. Therefore, for the KL calculation we obtained the mean (μ ~ (0, 1)) and covariance matrices (Σ ~(0, 1)) of the two *m*-dimensional distributions (N_0_, N_k_) and used them to calculate the KL divergence through:

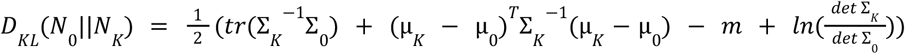

Reconstruction cost is also applied to model training as described in the original LFADS paper.

We used the Adam optimizer with gradient clipping to optimize the total alignment training loss. Total loss is obtained by a weighted sum of the above KL cost and reconstruction cost. During the training the learning rate was annealed, i.e., it was decreased through multiplication by a constant factor of 0.95 every time there was no improvement in the validation loss for a certain number of training epochs. The training procedure stops when validation loss has not shown any improvement for a fixed number of consecutive epochs. We selected the model weights corresponding to the lowest validation loss as the final model weights for inference.

Hyperparameters used for NoMAD Day K training are shown in Table 4.

**Table 4.**
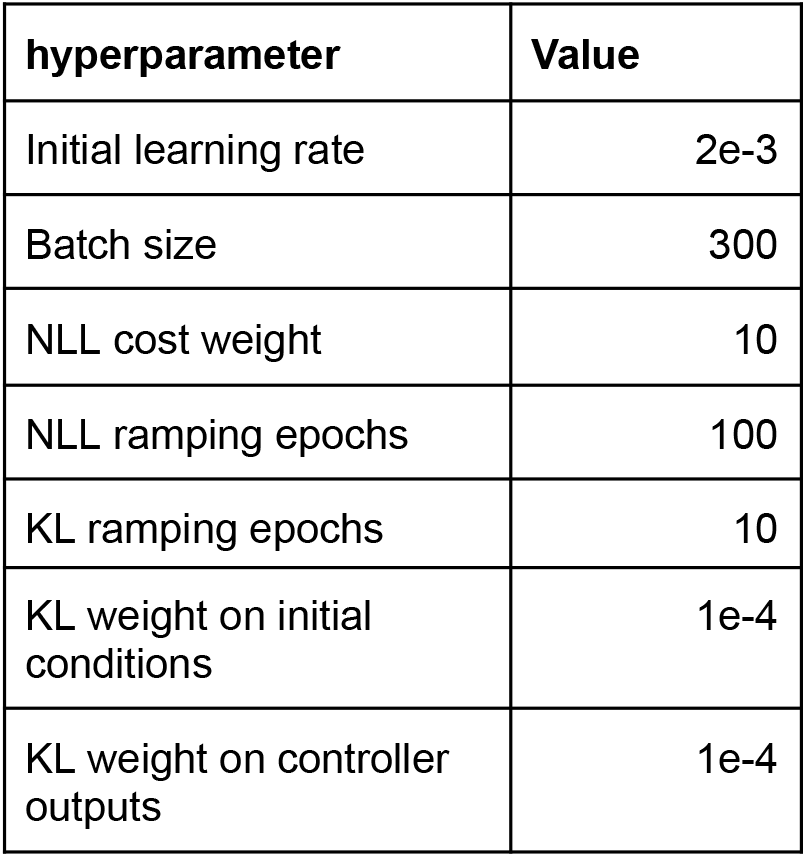
NoMAD hyperparameters.

**Table 5.** Aligned FA hyperparameters.

| parameter | value |
| --- | --- |
| Latent dimensionality | 10 |
| Number of FA models to fit | 5 |
| Maximum number of EM iterations to fit FA | 100000 |
| EM stopping criteria | 0.00001 |
| Minimum private variance threshold | 0.1 |
| Number of rows of loading matrix to use for alignment | 90 |
| Alignment L2 norm threshold | 0.01 |

### Comparisons

#### Degenhart et al., 2020 Aligned Factor Analysis (FA) Approach

The Degenhart et al. algorithm uses the following high-level procedure. First, the method fits a “Baseline Stabilizer” on the initial data, retaining some number of latents. This relies on factor analysis, which is not guaranteed to converge to an optimal representation. Thus their approach fits multiple FA models with random initialization, and they select the model with the highest log-likelihood. After fitting the baseline stabilizer, they also fit the data to be aligned with an FA model (same procedure as step 1). Next, they identify stable loading rows between the two models. This consists of iteratively trying to align the two loading matrices. After each alignment, they identify rows that are the most different after each alignment, and remove them. Finally, they learn the optimal orthonormal transformation to align the identified stable rows (i.e., solving the “Procrustes problem”). For our comparisons, we used the following parameters:

Based on data presented in Ma, et al. 2022, smoothing binned spike data prior to alignment with this approach improves performance^19^. Therefore, we use 20ms binned spike data smoothed with a 40ms Gaussian kernel as input data to this method. Only data from within behavioral trials is used to train this method, and behavioral trials containing outliers as determined in *Behavioral Outlier Removal* were discarded.

#### Adversarial Domain Adaptation Network (ADAN)

This method begins by fitting an autoencoder to reproduce smoothed binned spiking data and an RNN decoder to predict force (isometric monkey) or cursor velocity (kinematic monkey) activity from the manifold. To ensure a good Day 0 fit, we train this autoencoder using 5-fold cross validation and select the model with the best force R^2^. Then, ADAN is trained in a method similar to that of generative adversarial networks (GANs). A discriminator network is an autoencoder that acts to maximize the difference between the neural reconstruction losses on the two days. The distribution alignment module (the generator) works against the discriminator by minimizing the neural reconstruction losses on Day K. This results in alignment of the Day K manifold to the Day 0 manifold. We use the following parameters detailed in Table 6 to train this method:

**Table 6.** ADAN hyperparameters.

| parameter | value |
| --- | --- |
| Decoder latent dimensionality | 10 |
| Decoder batch size | 64 |
| Decoder epochs | 400 |
| Decoder learning rate | 1e-3 |
| Decoder steps | 4 |
| Decoder layers | 1 |
| ADAN epochs | 200 |
| ADAN batch size | 4 |
| ADAN discriminator learning rate | 5e-5 |
| ADAN generator learning rate | 1e-4 |

As input data to this method, we use 20ms binned spiking data smoothed with a 40ms Gaussian kernel. Only data from within behavioral trials is used to train this method and behavioral trials with outliers are discarded (see *Behavioral Outlier Removal*).

#### Static Decoder

Binned spikes (20ms bins) were smoothed with a Gaussian kernel (40ms). A decoder was trained on the Day 0 smoothed spikes. This fixed decoder was applied to the Day K smoothed spikes and evaluated.

### Neural Decoding

#### Wiener Filter

For both the kinematic and isometric datasets, prediction of behavioral output was done using a Wiener filter. Wiener filters predict the current value of an output signal using previous timesteps, as defined by:

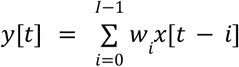

where *y[t]* is the output signal at time *t, x[t]* is the input signal at time *t*, *w_i_* is the filter coefficient, and *I* is the number of previous samples to use for decoding. In our decoder, the input signal *x* is the (aligned) manifold, *y* is the behavioral output to predict, and *I* was set to 4 time bins of history. The weights are fit using a matrix formation of the above equation:

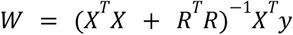

where *W* is a matrix of filter coefficients, *X* represents the predictor data with history and bias, and *y* represents the output signal. *R* represents a diagonal matrix with the L2 regularization constant filling the diagonal. The bias term is not regularized and therefore its diagonal entry is set to zero.

The L2 regularization aims to avoid decoder overfitting by penalizing solutions with large individual weights. L2 regularization values are obtained using 10-fold cross validation. We sweep a range of 20 values spanning 1e1 to 1e5 in logspace. For each value, we train and test a Wiener filter using 10-fold cross validation, testing the decoder on a held-out fold. The optimal regularization value was selected based on which value yielded the highest performance metric. Final performance was reported on the held out fold.

As input to the Wiener Filter, we use the generator states (*x*) and behavior (*y*). Each trial was represented as a window 250ms before to 500ms after movement onset. For each trial, the movement onset point was calculated using the period 250ms before the go cue to 750ms after the go cue. We first searched within this time period to identify the point at which the cursor reaches its maximum speed. From that point, we searched backwards in time to identify the point at which the cursor last reached 20% of its maximum speed. In parallel, we searched forward in time, beginning at the go cue, to find the point at which the cursor first reached 20% of its maximum speed. These two points should be consistent—if not, it indicates a trial in which the monkey started a movement, stopped, and then started again. We rejected trials with inconsistent movement onset calculations. We further rejected trials in which the backward move onset (last time the cursor reaches 20% of max speed) occurred before the point at which the target is displayed—this typically indicated that the monkey had not yet begun its movement in the time period analyzed, potentially due to inattention. While these trial rejections have minor effects on the analysis, we performed them to ensure the decoding metrics were a consistent and robust indicator of each method’s performance.

Only trials for which the monkey successfully completed the trial and movement onset was successfully calculated were considered. To account for drifts in the behavioral data baseline prior to movement onset that occurred across days, we manually set the first point in each trial’s force or velocity to zero and shift the remaining points accordingly.

#### Evaluation of Decoding Performance

The accuracy of neural decoding was measured using R^2^, defined as:

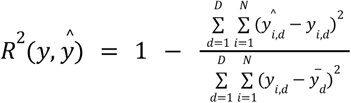

where *D* is the number of dimensions of the predicted output, *N* is the number of data samples, *y_i,d_* is an actual data sample for one dimension, 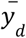 is the mean of the actual signal in one dimension, and 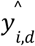 is a predicted data sample for one dimension. In practice, we used the function `sklearn.metrics.r2 score (y, y_hat, multioutput=‘variance_weighted’`) ^49^.

### Manifold Visualizations

In order to create visualizations of the manifold at different stages of the alignment process, we used demixed principal components analysis (dPCA)^36^. We applied regularized dPCA on the Day 0 manifold. We restricted dPCA fitting to successful trials within the window 250 ms before to 500 ms after target onset. After learning the Day 0 dPCA transformation, we applied the same transformation to the Day K manifold using both the Day 0 dPCA weights and the Day 0 mean offsets.

Visualizations were created by plotting the top condition-independent components and the top two condition-dependent components, as ranked by variance explained. This allows for the comparison of Day 0 to Day K before and after alignment without dependence on neural decoding or behavior.

### Calculating Decline in Decoding Performance Over Months

To quantify how much the decoding performance declined over the available timespan, we computed the rate of decay of the median performance within each 5-day bin. In order to reduce the effect of highly negative data points, we first convert R^2^ to a signal-to-noise ratio (SNR) metric using the equation *SNR* : =− 10 *log*_10_(1 – *R*^2^) as described in Makin et al., 2018^50^. We then fit an exponential decay equation (*y = Ae^−Bt^*) to the data where *y* is the median SNR within each 5-day bin with the median value of the Day 0 within-day performance appended as the first data point, and *t* is the middle of each bin with *t=0* appended as the first data point. This fits the constants *A* and *B* where *B* is the decay constant in units of 1/days. We convert this to half life in days for ease of interpretation using 1/*B* * *ln*(2).

## Supplementary Figures

**Supplementary Fig. 1.**
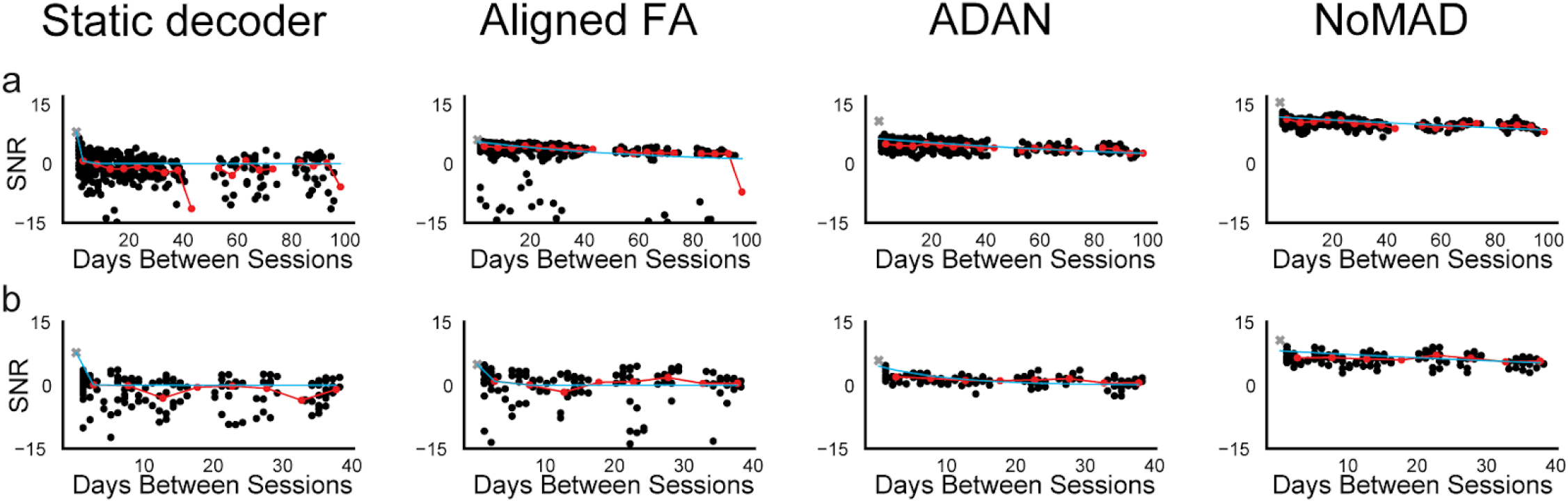
Exponential decay of decoding performance by method. Exponential curve fits (blue) to the median data (red) for (a) the isometric force datasets and (b) the reaching datasets. Decoding performance (R^2^) for individual pairs of days is shown by the black dots. The median R^2^ of the Day 0 decoder or classifier applied within the training day is shown by the gray x.

